# Integrated Multi-Omics Analyses of Synaptosomes Revealed Synapse-Centered Novel Targets in Alzheimer’s Disease

**DOI:** 10.1101/2025.01.09.631584

**Authors:** Subodh Kumar, Enrique Ramos, Axel Hidalgo, Daniela Rodarte, Bhupender Sharma, Melissa M. Torres, Davin Devara, Shrikanth S. Gadad

## Abstract

Synapse dysfunction is an early event in Alzheimer’s disease (AD) caused by various factors such as Amyloid beta, p-tau, inflammation, and aging. However, the exact molecular mechanism of synapse dysfunction in AD is largely unknown. To understand this, we comprehensively analyzed the synaptosome fraction in postmortem brain samples from AD patients and cognitively normal individuals. We conducted high-throughput transcriptomic analyses to identify changes in microRNA (miRNA) and mRNA levels in synaptosomes extracted from the brains of both unaffected individuals and those with Alzheimer’s disease (AD). Additionally, we performed mass spectrometry analysis of synaptosomal proteins in the same sample group. These analyses revealed significant differences in the levels of miRNAs, mRNAs, and proteins between the groups. To further understand the pathways or molecules involved, we used an integrated omics approach and studied the molecular interactions of deregulated synapse miRNAs, mRNAs, and proteins in the samples from individuals with AD and the control group, which demonstrated the impact of deregulated miRNAs on their target mRNAs and proteins. Furthermore, the DIABLO analysis highlighted complex relationships between mRNAs, miRNAs, and proteins that could be key in understanding the pathophysiology of AD. Our study identified synapse-centered novel candidates that could be critical in restoring synapse dysfunction in AD.

## Introduction

Alzheimer’s disease (AD) is a neurodegenerative disorder that is responsible for 60-80% of dementia cases in the United States (Alzheimer’s Facts and Figures, 2023) (1). There is a myriad of factors that have been associated with AD, such as aging, genetics, lifestyle, infections, and injuries (2). They can contribute to neurodegeneration and cognitive decline in the patients. The two main pathological hallmarks of AD, amyloid-beta (Aβ) plaques and neurofibrillary tangles (NFTs) made of hyperphosphorylated tau (p-tau), are thought to be responsible for neural deterioration (3). Recent studies have shown that one of the earliest proponents of AD is synaptic dysfunction, which subsequently leads to synaptic degeneration (4–7). Both Aβ oligomers and NFTs have been implicated in synaptic dysfunction via altering multiple synaptic events (4–6,8).

Synapses are fundamental units that allow neuronal communication throughout the nervous system and with other brain cell types (9). Functioning synapses are essential for normal cognitive functions and brain activities. As such, it is not surprising that the synaptic dysfunction in AD presents with cognitive deficiencies. Aβ-mediated synaptic dysfunction is one of the heavily studied mechanisms of AD pathology. In 1998, Aβ oligomers were found to cause the loss of dendritic spines, often the postsynaptic component (10). This disrupts NMDA-dependent long-term potentiation (LTP) and instead facilitates long-term depression (LTD) (11). Further studies also found Aβ in the presynaptic terminals of glutamatergic neurons, in which they upregulate the amount of glutamate in the synaptic cleft. This causes excitotoxicity and desensitization of glutamate receptors, contributing to synaptic dysfunction (11). Meanwhile, p-tau is found in both presynaptic and postsynaptic terminals and induces synaptic dysfunction through its own mechanisms. These include impairing presynaptic vesicle release, maturation of dendritic spines, and mitochondrial function in synapses. P-tau can also trigger synaptic phagocytosis by microglia (8). While we understand that Aβ plaques, p-tau, aging, and inflammation all play roles in synaptic dysfunction, the exact molecular mechanism in which it occurs is still unknown (5,12,13). Recently, synaptosomes have been used to study synaptic dysfunction at a molecular level in AD and other neurodegenerative diseases. They are intact forms of synapses, containing all the players involved in synaptic transmission. This includes the presynaptic and postsynaptic membranes, mitochondria, proteins, neurotransmitters, mRNA, microRNA (miRNA), and more (5,15,16).

One of the components of synaptic dysfunction that has been gaining interest is miRNAs. They are small, non-coding RNAs that regulate gene expression. In regards to AD, they have been implicated in modifying synaptic protein expression and transcription factors that lead to synaptic dysfunction (5,15,17–20). Many studies have identified specific miRNAs that play crucial roles in the synapses of AD, such as miR-34a, miR-92, miR-125b, and much more (18,21–23). In 2022, our lab also identified miR-501-3p, miR-502-3p, and miR-877-5p as novel synaptosomal miRNAs upregulated with AD progression (5). Since many different miRNAs are differentially expressed in AD, they have been heavily studied as potential biomarkers (18,24–28). While many studies have explored the mechanisms of miRNA-related pathologies in AD, we are still unsure how specific synaptosomal molecules change in the setting of synaptic dysfunction.

Dysregulation of mRNA expression and its proceeding protein translation in AD has also been studied. Hundreds of differentially expressed mRNAs have been identified in AD. Among them, genes affecting the function of microglia and astrocytes were shown to be associated with a clinical diagnosis of AD (29). Genes affecting the electron transport chain and protein binding were also involved in AD (30). While identifying differentially expressed genes is important, mRNA expression only accounts for 40% of protein variance in mammals (31). Proteomic studies are necessary to convey a more complete understanding of AD pathology. Aside from Aβ and tau, several other synaptic proteins have been identified in AD. These include Calsyntenin-1, GluR2, GluR4, and Neurexin-2A (32). These proteins have been investigated for their potential as synaptic AD biomarkers; however, little is known about how they directly contribute to synaptic dysfunction.

To this date, no study has been done to assess the interplay between synapse miRNAs, mRNAs, and proteins in the setting of synapse dysfunction in AD. It is unclear how a specific miRNA, mRNA, and protein change in the AD synapse relative to the control’s synapse. It is also unclear if the deregulation of all three molecular subsets is connected and if their expression is dependent on each other. Therefore, to understand the status of each molecular subset, here we conducted the transcriptomic and proteomic analysis of miRNA, mRNA, and protein in AD and controlled synapse. We also implemented a multi-omics integrative approach to assess how each component changes in AD and analyze their interactions to understand the molecular basis of synaptic dysfunction. The current study provides synapse-centered novel molecular signatures that could be potential therapeutic and synaptic biomarker targets in AD.

## Materials and Methods

### Postmortem brain samples

Postmortem brains from AD patients and unaffected controls were obtained from NIH NeuroBioBanks-(1) Human Brain and Spinal Fluid Resource Center, 11301 Wilshire Blvd (127A), Los Angeles, CA. (2) Brain Endowment Bank, University of Miami, Millar School of Medicine, 1951, NW 7th Avenue Suite 240, Miami, FL. (3) Mount Sinai NIH Brain and Tissue Repository, 130 West Kingsbridge Road Bronx, NY. Brain tissues were dissected from AD patients from Brodmann’s Area 10 of the frontal cortices (n = 27) and age and sex-matched unaffected controls (n = 14). Demographic and clinical details of study specimens are provided in **Supplementary Table 1**. The study was conducted at the Molecular and Translational Medicine Department, Texas Tech University Health Sciences Center El Paso, and the Institutional Biosafety Committee (IBC protocol # 22008) approved the study protocol for the use of human postmortem brain tissues obtained from NIH NeuroBioBanks. The NIH NeuroBioBanks mentioned above operated under their institution’s IRB approval, and they obtain written informed consent from the donors (5,33).

### Synaptosome extraction

Synaptosomes were extracted using Syn-PER Reagent as described in our previous study (5). Briefly, 50 mg of brain tissue was used from each sample for synaptosome extraction in 1 ml of Syn-PER Reagent. Tissues were homogenized slowly by Dounce glass homogenization on ice with ∼10 slow strokes. Samples were centrifuged at 1400 g for 10 minutes at 4 °C to remove the leftover tissue debris. After centrifugation, the supernatant was transferred to a new tube. Again, the supernatant (homogenate) was centrifuged at a speed of 15,000 g for 20 min at 4 °C. The supernatant was removed as a cytosolic fraction, and synaptosomes were recovered in pellet form. The synaptosome pellets were processed for RNA and protein extraction.

### MiRNA- and mRNA-Seq analysis

Total RNA, including miRNAs, was extracted from the synaptosomes of AD and control postmortem brain samples using the TriZol reagent with some modifications. RNA quality and purity were determined by NanoDrop. The mRNAs and miRNAs HiSeq analysis were performed commercially at LC Sciences Houston, Texas. MiRNA HiSeq analysis flow chart is shown in the Supplementary Information Figure 1 (**SI Figure 1**).

**Figure 1.**
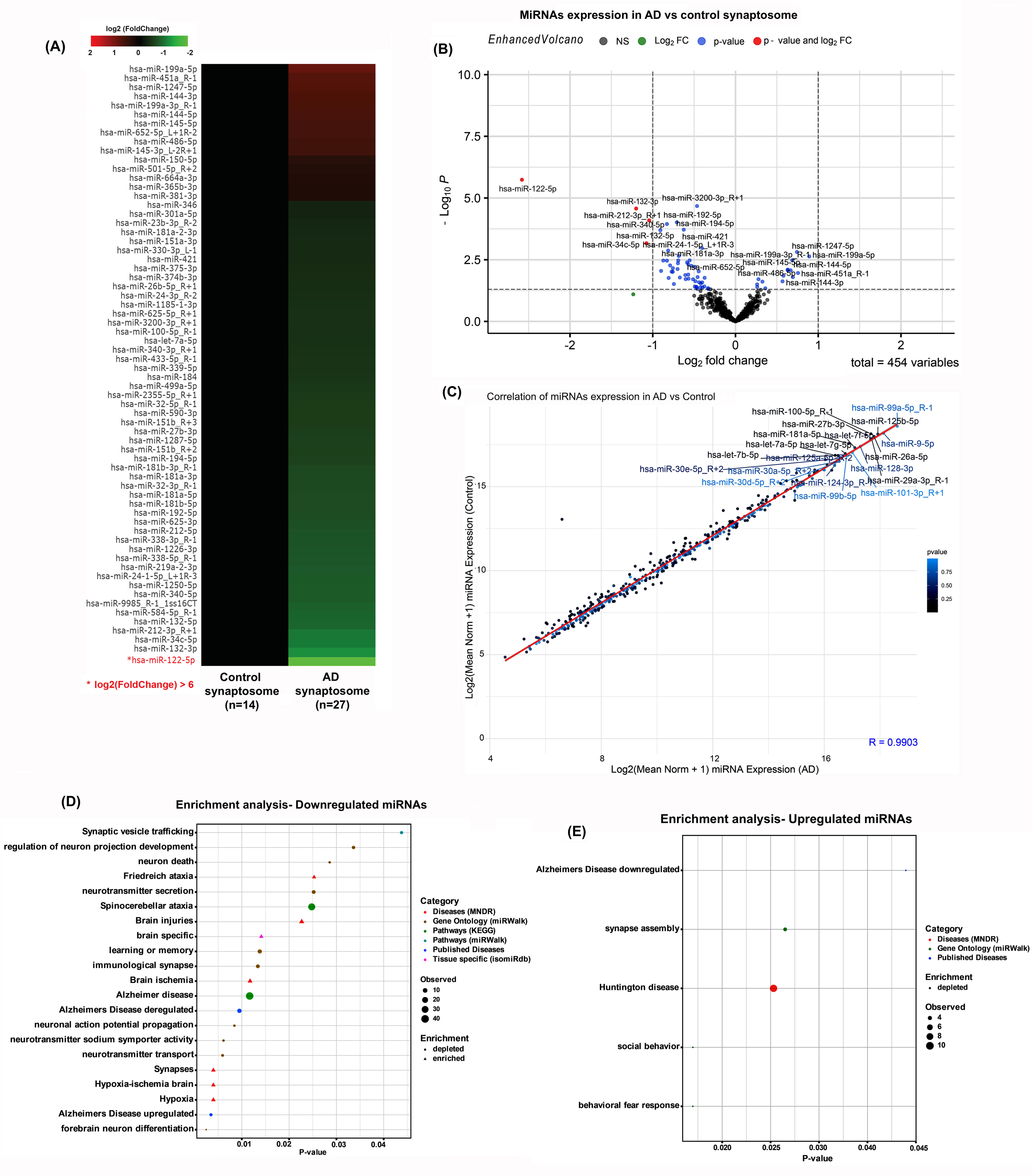
MiRNA-Seq analysis in AD vs control synapse. **(A)** Heatmap displaying differentially regulated miRNAs: the top upregulated and downregulated miRNAs in Alzheimer’s disease (n=27) compared to control (n=14) synaptosomes. **(B)** A volcano plot showing the top deregulated miRNAs’ (log_10_ p-value). **(C)** Correlation analysis of the top differentially regulated miRNAs in AD versus control synaptosomes with a significant R-value. **(D)** Gene set enrichment analysis of top downregulated miRNAs shows affected biological pathways in human diseases, along with their p-values and number of genes. **(E)** Gene set enrichment analysis of top-upregulated miRNAs shows depleted and enriched biological pathways in human diseases, along with their p-values and numbers of genes.

### RNA (polyA) sequencing

RNA-Seq (RNA sequencing), also called whole transcriptome shotgun sequencing (WTSS), uses next-generation sequencing (NGS) to reveal the presence and quantity of RNA in a biological sample at a given moment in time. RNA-Seq was used to analyze the continuously changing cellular transcriptome. Specifically, RNA-Seq facilitates looking at alternative gene spliced transcripts, post-transcriptional modifications, gene fusion, mutations/SNPs, and changes in gene expression over time or differences in gene expression in different groups or treatments. In addition to mRNA transcripts, RNA-Seq can look at different populations of RNA, including total RNA and small RNA, such as miRNA, tRNA, and ribosomal profiling. RNA-Seq can also determine exon/intron boundaries and verify or amend previously annotated 5’ and 3’ gene boundaries. Prior to RNA-Seq, gene expression studies were done with hybridization-based microarrays. Issues with microarrays include cross-hybridization artifacts, poor quantification of lowly and highly expressed genes, and the need to know the sequence a priori. Because of these technical issues, transcriptomics transitioned to sequencing-based methods. These progressed from Sanger sequencing of Expressed Sequence Tag libraries to chemical tag-based methods (e.g., serial analysis of gene expression) and finally to the current technology, next-gen sequencing of cDNA (notably RNA-Seq).

### Library construction and sequencing

Poly(A) RNA sequencing library was prepared following Illumina’s TruSeq-stranded-mRNA sample preparation protocol. RNA integrity was checked with Agilent Technologies 2100 Bioanalyzer. Poly(A) tail-containing mRNAs were purified using oligo-(dT) magnetic beads with two rounds of purification. After purification, poly(A) RNA was processed for DNA library construction. Quality control analysis and quantification of the sequencing library were performed using Agilent Technologies 2100 Bioanalyzer High Sensitivity DNA Chip. Paired-ended sequencing was performed on Illumina’s NovaSeq 6000 sequencing system.

All extracted RNA was used in the library preparation following Illumina’s TruSeq -small-RNA-sample preparation protocols (Illumina, San Diego, CA, USA). Quality control analysis and quantification of the DNA library were performed using Agilent Technologies 2100 Bioanalyzer High Sensitivity DNA Chip. Single-end sequencing 50bp was performed on Illumina’s Hiseq 2500 sequencing system following the manufacturer’s recommended protocols.

### Bioinformatics analysis

#### miRNA

Raw reads were subjected to an in-house program, ACGT101-miR (LC Sciences, Houston, Texas, USA) to remove adapter dimers, junk, low complexity, common RNA families (rRNA, tRNA, snRNA, snoRNA) and repeats. Subsequently, unique sequences with length in 18∼26 nucleotide were mapped to specific species precursors in miRBase 22.0 by BLAST search to identify known miRNAs and novel 3p- and 5p-derived miRNAs. Length variation at 3’ and 5’ ends and one mismatch inside the sequence was allowed in the alignment. The unique sequences mapping to specific species of mature miRNAs in hairpin arms were identified as known miRNAs. The unique sequences mapping to the other arm of known specific species precursor hairpin opposite to the annotated mature miRNA-containing arm were considered novel 5p- or 3pderived miRNA candidates. The remaining sequences were mapped to other selected species precursors (with the exclusion of specific species) in miRBase 22.0 by BLAST search, and the mapped pre-miRNAs were further BLASTed against the specific species genomes to determine their genomic locations. The unmapped sequences were BLASTed against the specific genomes, and the hairpin RNA structures containing sequences were predicated from the flank 80 nt sequences using RNAfold software (http://rna.tbi.univie.ac<u>. at/cgi-bin/RNAfold.cgi</u>). The criteria for secondary structure prediction were: (1) number of nucleotides in one bulge in the stem (≤12), (2) number of base pairs in the stem region of the predicted hairpin (≥16), (3) cutoff of free energy (kCal/mol ≤-15), (4) length of hairpin (up and down stems + terminal loop ≥50), (5) length of hairpin loop (≤20), (6) number of nucleotides in one bulge in mature region (≤8), (7) number of biased errors in one bulge in mature region (≤4), (8) number of biased bulges in mature region (≤2), (9) number of errors in mature region (≤7), (10) number of base pairs in the mature region of the predicted hairpin (≥12), (11) percent of mature in stem (≥80).

### Analysis of differentially expressed miRNAs

Differential expression of miRNAs based on normalized deep-sequencing counts was analyzed selectively using the Fisher exact test, Chi-squared 2X2 test, Chi-squared nXn test, Student t test, or ANOVA, depending on the experiment’s design. The significance threshold was set to 0.01 and 0.05 in each test.

### The prediction of target genes of miRNAs

To predict the genes targeted by the most abundant miRNAs, two computational target prediction algorithms (TargetScan 50 and Miranda 3.3a) were used to identify miRNA binding sites. Finally, the data predicted by both algorithms were combined, and the overlaps were calculated. The GO terms and KEGG Pathway of these most abundant miRNAs and miRNA targets were also annotated.

#### <u>mRNA</u>, Transcriptome assembly and differential expression

Sequencing and mapping of NGS paired-end reads (150bp) to the human transcriptome (hg38) with annotation from Ensembl release version 101 (August 2020) using the HISAT2 aligner (34). SAM files were converted into BAM files and sorted and indexed using Samtools (35). Read counts were computed using the alignment files and the FeatureCounts program from the subread-2.0.1 package (36) with reference annotation from Ensembl version 101. Counts normalization and differential gene expression analyses were calculated using the DESeq2 R package (37). Filtering criteria for differentially expressed genes was set to P-adjusted value < 0.05. The data was visualized using R statistical software, including packages: ggplot2, plotly, pheatmap, and enhancedVolcano.

### Gene Ontology and KEGG Pathways

Biotype classification and composition analyses were done using the Ensembl database (REST API) on filtered differentially expressed genes. Gene Ontology (GO) was carried out by querying the bioinformatic Database for Annotation, Visualization, and Integrated Discovery (DAVID) (38). The following annotation categories were included in the analysis: KEGG Pathways, GO: Biological Processes, GO: Molecular Function, GO: Cellular Components. The data was visualized in R statistical software (version 4.0.3) using the packages ggplot2 and plotly/heatmap for dot plots and heatmaps, respectively.

#### <u>Protein,</u> Mass spectrometry analysis

The LC-MS Mass spec analysis of synaptosome proteins was conducted in 5 AD and 5 control samples with technical duplication using the Mass Spectrometry Research Facility, University of South Alabama, Alabama.

### Sample preparation

The synaptosomal proteins from AD and control postmortem brain tissues were prepared by using the SynPer kit from Thermofisher, followed by protein digestion. Pellets were stored at - 80°C upon receipt and thawed at room temperature immediately before preparation. 25μl 8M urea was added to the pellets and mixed at 37°C for 10 minutes at 600 rpm. The samples were diluted with 200μl 50mM ammonium bicarbonate (ABC) containing 10 mM tris-carboxyethyl phosphine (TCEP) for reduction. In-solution protein digestion was carried out using 4μl sequencing-grade modified porcine trypsin (0.8μg) (Promega, Madison, WI) overnight at 37°C in a shaker at 600 rpm. Centrifuged samples at 16,100 × g for 15 min at 4 °C in the tabletop microcentrifuge. Transferred 200 μl of the supernatant to a snap–top autosampler vial for HPLC isolation.

### Mass Spectrometry Data Analysis

In this study, mass spectrometry (MS) data were analyzed using a combination of MaxQuant software for peptide and protein identification and quantification, followed by statistical analysis in R using the Proteus package (39–41). Initially, the raw MS data files generated by the liquid chromatography-tandem mass spectrometry (LC-MS/MS) experiment were imported into MaxQuant (version 2.6.5). Key parameters in MaxQuant were set to include a 1% false discovery rate (FDR) for both peptide and protein identification, ensuring high-confidence results. The search was conducted against a species-specific protein database, with parameters set for enzyme specificity (e.g., trypsin), a defined number of missed cleavages, and variable and fixed modifications, such as oxidation of methionine and carbamidomethylation of cysteine, respectively.

Label-free quantification (LFQ) was used to assess protein abundance across samples without the need for stable isotope labeling. The “match between runs” feature was enabled, which helps to match peptide features across different LC-MS/MS runs, thus increasing the number of identified peptides and improving protein quantification consistency across replicates. After running the MaxQuant pipeline, the results, including the *proteinGroups.txt* file, were exported for further analysis.

Next, the output from MaxQuant was loaded into R for analysis using the Proteus R package. Proteus offers a robust framework for proteomics data analysis, including functions for filtering low-quality data, normalization, and statistical testing. Initially, the data were filtered to remove proteins with poor identification or low coverage, ensuring the analysis was focused on high-confidence protein identifications. LFQ intensity values were then normalized to correct for technical variation across samples. Statistical testing for differential protein expression between conditions was performed using moderated t-tests or other appropriate methods depending on the experimental design, with Proteus handling the multiple hypothesis testing correction using the

Benjamini-Hochberg procedure to control for false positives. The results included lists of significantly differentially expressed proteins, which were further interpreted using downstream bioinformatics tools for functional enrichment and pathway analysis (39–41).

### Western blot analysis

The proteins were extracted from the AD and control postmortem brains. Briefly, 20 mg brain tissues were suspended in RIPA buffer (Thermo Scientific) supplemented with protease inhibitors (Thermo Scientific) and disrupted by ultra-sonication (Qsonica, USA; amplitude 80%, pulse 10 sec on/ off, time 30 seconds). Thereafter, tissue debris were removed by centrifugation (13,000 rpm for 20 minutes). Protein concentration was estimated by BCA assay (Thermo Scientific). Equal amounts of protein (40 μg per sample) were separated by SDS-PAGE on 10% polyacrylamide gels and transferred onto PVDF membranes (BioRad, Hercules, CA, USA). Membranes were blocked in 5% BSA for 1 hour at room temperature. The membranes were then incubated overnight at 4°C with primary antibodies against GPI, UQCRC1, TIMM50, and VAT1L and GAPDH (1:1000 v/v, Proteintech). Details of the antibodies and their dilutions used are listed in **Supplementary Table 2**. After washing the membranes three times with TBS-T buffer at 10-minute intervals, they were incubated for 1 hour at room temperature with a secondary antibody (rabbit anti-mouse horseradish peroxidase [HRP] 1:10,000). Following three additional washes with TBS-T buffer, proteins were detected using chemiluminescence reagents (Thermo Fisher Scientific) and visualized with an Amersham imager 680 (GE Healthcare Bio-Sciences, Uppsala, Sweden). Protein band intensities were quantified using ImageJ software (1.54d, Java 1.8.0_345: http://imagej.org) for densitometry analysis as described in more detail in our earlier study (42), and relative protein expression levels were normalized to GAPDH as loading controls.

### Multi-omics integration analysis

Identifying genes with high variance: Identifying genes with high variance across biological replicates involves detecting genes whose expression levels exhibit significant variability among different samples of the same condition. High variance can indicate biologically relevant differences or technical noise affecting gene expression consistency. We conducted a comprehensive analysis using the R programming language to identify these genes, leveraging the capabilities of the “limma” and “edgeR” packages.

### Data Preparation

Creation of DGEList Object: We started by creating a DGEList object from the raw count data, which includes the counts of reads mapped to each gene across all samples. This object is essential for downstream analysis in edgeR. Filtering Lowly Expressed Genes: To ensure the robustness of our analysis, we filtered out genes with low expression levels across samples. This step reduces noise and improves the detection of genuinely variable genes.

### Normalization and Transformation

Normalization of Counts: We normalized the raw counts for differences in sequencing depth and RNA composition using the trimmed mean of M-values (TMM) method in edgeR. Normalization ensures that comparisons of gene expression levels across samples are accurate. Variance-Stabilizing Transformation: We applied the voom transformation from the limma package to stabilize the variance across the mean expression levels. This transformation is crucial for making the data suitable for linear modeling.

### Linear Modeling and Statistical Analysis

Fitting the Linear Model: Using the lmFit function from the limma package, we fit a linear model to the transformed data. This step involves estimating the expression levels of genes and accounting for the experimental design. Computing Statistics with eBayes: We employed the eBayes function to compute empirical Bayes statistics for the linear model coefficients. This approach improves the reliability of statistical inferences by borrowing strength from the ensemble of genes. Extracting Top Genes by P-Value: Finally, we ranked the top genes with high variance based on their P-values. Genes with the lowest P-values were considered the most significant variable across replicates.

Using edgeR and limma in our analysis provided a robust framework for identifying genes with high variance in expression levels across biological replicates. By combining normalization, variance-stabilizing transformation, linear modeling, and empirical Bayes statistics, we ensured the identification of genes exhibiting genuine biological variability, minimizing the impact of technical noise.

### Statistical analysis

The statistical analyses of biological data were conducted using the student’s *t*-test to analyze two groups of samples: control *versus* AD miRNAs, mRNAs, and protein analysis. One-way variance analysis was used to analyze the data between three groups of samples, such as synapse protein changes across the controls and AD samples with different Braak stages. Statistical parameters were calculated using GraphPad Prism software, v6 (GraphPad, San Diego, CA, USA) (www.graphpad.com). The *P* values < 0.05 were considered statistically significant.

### DIABLO Analysis

We performed a multi-omics integration analysis using the DIABLO (Data Integration Analysis for Biomarker discovery using Latent cOmponents) framework available in the mixOmics package in R programming language (43). The analysis involved the following steps: Sample Selection: Only 5 samples per group (AD and HC) had proteomics data, so we only included those samples for further analysis for mRNA-seq, miRNA-seq, and proteomics. *Data Integration:* We combined the preprocessed datasets into a list format suitable for DIABLO analysis. Each omics dataset was represented as a data frame where rows corresponded to samples and columns to features (genes, miRNAs, or proteins). *Design Matrix Specification:* To facilitate the integration of these datasets, we defined a design matrix that specified the connection strength between each pair of datasets. The diagonal elements of the design matrix were set to 0, while off-diagonal elements were set to 0.1 to allow for moderate integration strength. *Model Fitting:* We set the number of components to 5, allowing the model to capture key latent variables across the datasets. The DIABLO model was fitted to the data using the “block.splsda” function from the mixOmics package. *Visualization and Analysis:* We visualized the sample projections on the latent components using the “plotIndiv” function, which allows clustering patterns to be examined according to the outcome variable. The contribution of variables to each component was visualized using the “plotVar” function. A Heatmap of the integration was carried out using the function “cimDiablo,” and a circos plot of the integration was done using the “circosPlot” function. *Extraction of Loadings:* To identify the key biomarkers, we extracted the loadings of each variable onto the components. We selected components 1 and 2 for such analysis. *Model Validation:* The performance and stability of the DIABLO model were assessed through cross-validation. We used repeated k-fold cross-validation to ensure the robustness of the selected components and the identified biomarkers.

## Results

### MiRNAs HiSeq analysis of synaptosomal miRNA in Alzheimer’s brain

The miRNA-Seq analysis was conducted on synaptosomal RNAs isolated from the cortical area (BA10) of 14 controls and 27 AD postmortem brains. A total of 965 precursor-miRNAs (pre-miRNA), 1440 mature miRNAs, and 486 novel Potential Candidates (PCs) were found to be expressed in the AD and control synapses (**Supplementary Table 3**). Among all categories of small RNAs, 1793 molecules, including pre-miRNAs, miRNAs, and PCs, were significantly deregulated in AD versus control synapses (**Supplementary Table 4**). **SI Figure 2** shows the complete heat map of all deregulated miRNAs in AD versus control synapses. Next, we sorted out only mature miRNA list using adjusted p-values and higher fold change. As a result, 115 mature miRNAs were found to be significantly deregulated (p < 0.05; fold change +/− 2) in AD versus control synapses (**Supplementary Table 5**). Further, to identify the topmost deregulated miRNAs, we narrowed down the miRNA selection criteria with miRNA reads (>10 per sample) and fold change (+/− 6-fold; p < 0.05), and we found 38 miRNAs that were more significantly deregulated in AD synapse relative to control synapse. The eight miRNAs were upregulated, and 30 miRNAs were found to be downregulated, as shown in the miRNA heatmap (**Figure 1A**). These miRNAs were further segregated based on their log2 fold change and p-values, as shown in the volcano plot (**Figure 1B**). MiR-122-5p, miR-132-3p, miR-212-3p, and miR-34c-5p showed the highest Log10 p-value and fold change variance. Further, miRNA correlation analysis was performed based on the log2 mean values in both AD and controls. Six miRNAs highlighted in blue displayed a significant correlation regarding their mean value in AD and controls, as shown in the correlation plot (**Figure 1C**). Further, we conducted the in-silico bioinformatic analysis of downregulated and upregulated miRNAs in AD synapses. The KEGG/GO enrichment analysis showed that downregulated miRNAs are involved in several brain and neuron-related cellular pathways (**Figure 1D**; **Supplementary Table 6**). Importantly, miRNAs are associated with AD, synapse, learning and memory, and other AD-related pathways. Similarly, the upregulated miRNAs were also involved in AD, synapse assembly, Huntington’s disease, social behavior, and behavioral fear response (**Figure 1E**; **Supplementary Table 7**). These results confirm that the deregulation of synapse miRNAs in AD is detrimental to normal synapse function, brain activity, and AD.

### mRNA-Seq analysis of synaptosomal mRNA in Alzheimer’s brain

mRNA or polyA RNA-Seq analysis was conducted on synaptosome mRNAs in the same 14 controls and 27 AD postmortem brains. A total of 937 mRNAs were found to be significantly deregulated (p<0.05; fold-change +/− 2) in AD versus control synaptosomes **(Supplementary Table 8)**. **SI Figure 3** showed the complete heat map of significantly deregulated mRNAs in AD versus control synapses. To identify the topmost deregulated mRNAs, we narrowed down the miRNA selection criteria with mRNA reads (>10 per sample), and fold change (+/−2) showed the top 20 significantly deregulated genes **(Figure 2A).** The heatmap showed the top ten upregulated genes were-*SHANK1*, *HIVEP3*, *SLC7A2*, *SAMD4A*, *BCL9L*, *GAREM2*, *GLIS3*, *MTSS2*, *NWD1*, and *KDM6B*. While downregulated genes were-*TF*, *TUBA1A*, *MAL*, *EIF5B*, *SEPTIN4*, *CNP*, *S100B*, *SEMA3B*, *MOBP* and *PLP1* **(Figure 2A)**. The same top gene transcripts were segregated in AD and controls based on their log2 fold change and p-values, as shown in the volcano plot **(Figure 2B)**. Next, correlation analysis exhibited a significant correlation between potential gene targets in AD versus controls in terms of their mean value in both AD and controls, as shown in the correlation plot **(Figure 2C)**. Further, we conducted the *in-silico* bioinformatic analysis to determine the roles of deregulated miRNAs in human disease, biological pathways, and cellular processes **(SI Figures 4 and 5)**. A KEGG/GO analysis of the upregulated mRNAs shows that most of these genes are involved in AD-related processes: PI3K-Akt signaling pathway, Ras signaling pathway, axon guidance, Rap1 signaling pathway, Wnt signaling pathway, MAPK signaling pathway, JAK-STAT signaling pathway, cAMP signaling pathway, mTOR signaling pathway, HIF-1 signaling pathway, TGF-beta signaling pathway, Hippo signaling pathway, FoxO signaling pathway, Hedgehog signaling pathway, and the Longevity regulating pathway **(SI Figure 4).** KEGG/GO analysis of the downregulated mRNAs also shows their involvement in AD-related cellular processes: metabolic pathways, glutathione metabolism, and cysteine and methionine metabolism **(SI Figure 5).** Additionally, we conducted the gene set enrichment analysis for the deregulated genes separately in controls and AD. The deregulated mRNA involved in multiple normal cellular process and biological pathways in healthy controls (**SI Figure 6**), while in AD cases deregulated mRNAs are mostly involved in neurological and synapse related biological pathways as shown by KEGG pathway analysis (**SI Figure 7**). These results confirmed that synaptosome-localized mRNA population levels are significantly altered in AD and associated with multiple neuronal pathways.

**Figure 2.**
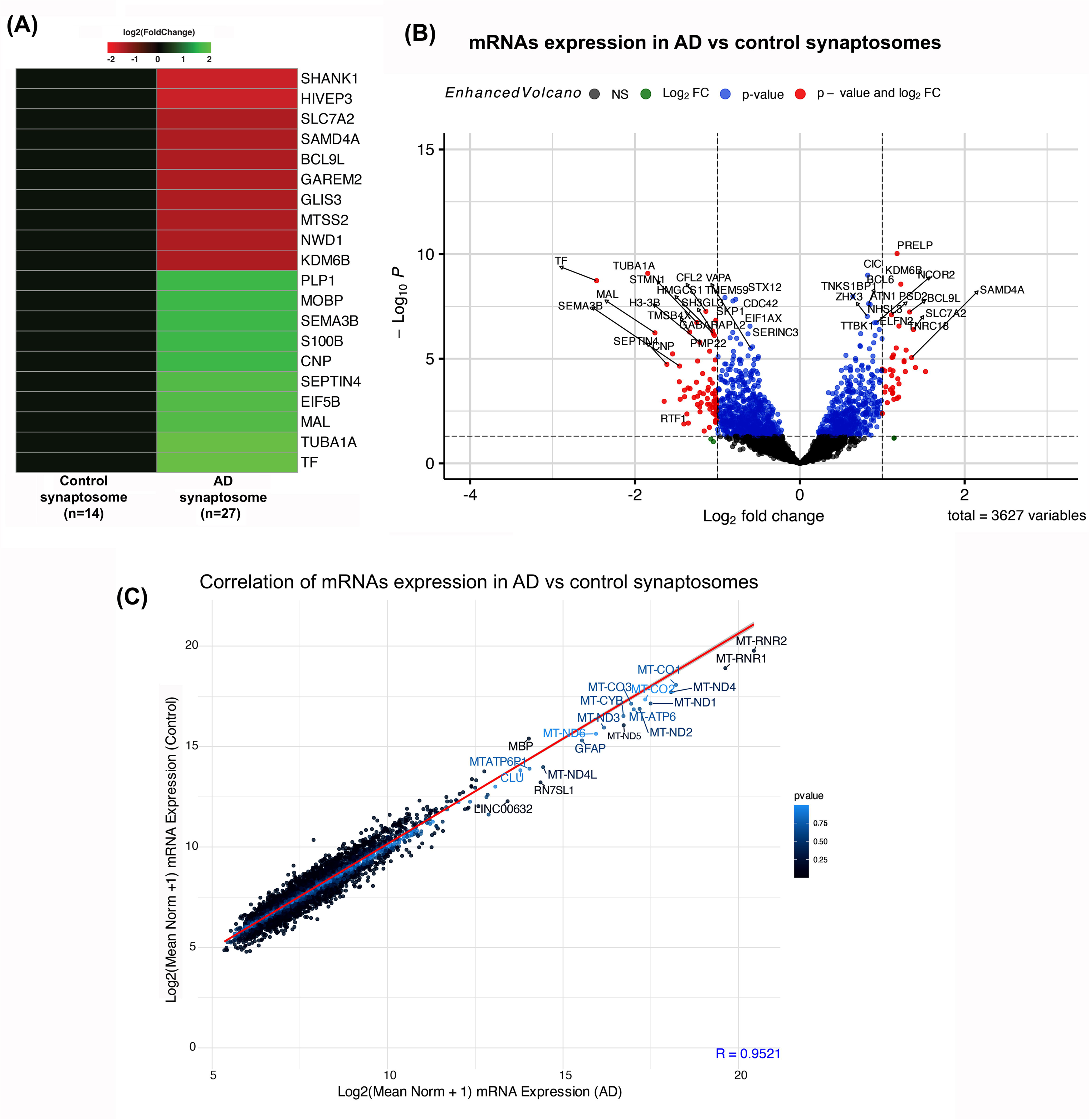
mRNA-Seq analysis in AD vs control synapse. **(A)** Heatmap showing the top differentially regulated genes in AD (n=27) versus control (n=14) synaptosomes. **(B)** Volcano plot depicting the top differentially regulated genes in AD versus control synaptosomes. **(C)** Correlation analysis of top genes in AD vs. control synaptosome with significant R-value.

### Mass spec analysis of synaptosomal proteins in Alzheimer’s brain

To identify differentially regulated proteins, we performed mass spec analysis on synaptosomal proteins from 5 controls and 5 AD postmortem brains (Braak stage VI) with two technical duplicates. A total of 1066 proteins were identified in the synaptosomal fraction in AD and controls (**Supplementary Table 9**). Furthermore, proteins were categorized based on their significant fold changes. A total of 152 proteins were determined to be significantly deregulated (p < 0.05; fold change +/− 2) in AD versus control synaptosomes (**Supplementary Table 10**). To pinpoint the most deregulated proteins, we narrowed down the protein selection criteria by using a fold change of (+3/−3) and p-values (p < 0.05). Seventeen proteins were identified as the most significantly upregulated (TIMM50, NONO, VAT1L, MOBP, ENPP6, PPP3R1, RDX, TF, HSPA2, VGF, H1-2, SEPTIN4, NEFM, MOG, RPLP2, INA, and BCL2L13), and sixteen proteins were found to be significantly downregulated (GPI, UQCRC1, DNAJA1, PGAM5, CASKIN1, FGG, CLIC4, VPS50, FGB, EEF1A2, SCRN1, GATD3, PYGM, YWHAE, YWHAG, and UCHL1) as shown in the protein heatmap (**Figure 3A**). The top deregulated proteins were further categorized based on their log2 fold change and Log10 p-values, as depicted in the volcano plot (**Figure 3B**). Several proteins exhibited a significant correlation regarding their Log2 mean value in AD and controls, as presented in the correlation plot (**Figure 3C**). Additionally, we performed in-silico bioinformatic analysis to determine the roles of deregulated proteins in human disease, biological process and cellular pathways. The KEGG pathway analysis of upregulated proteins revealed the significant involvement of upregulated synaptosomal proteins in key neurological pathways such as Pathways of neurodegeneration-multiple diseases, Parkinson’s disease, Long-term potentiation, and HIF-1 signaling pathway (**Figure 3D**). Similarly, upregulated proteins at synapses also participated in multiple biological pathways, including dopaminergic, cholinergic, and serotonergic synapses (**Figure 3E**). These findings confirmed that synaptosome-localized protein levels are significantly altered in AD.

**Figure 3.**
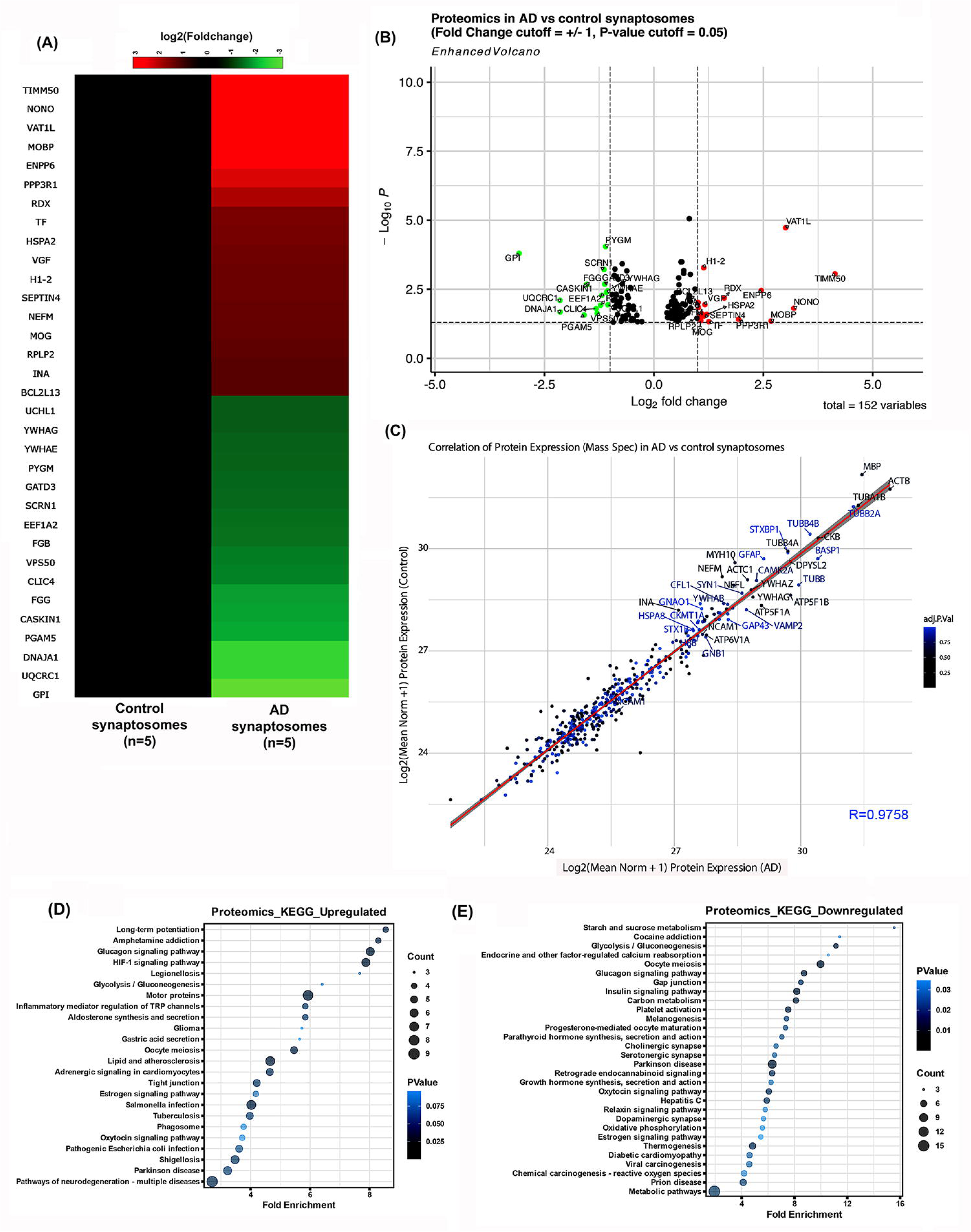
Mass-spec analyses of differentially regulated proteins in AD vs control synapse. **(A)** The heatmap of the top-up and downregulated proteins in AD (n=5) versus control (n=5) synaptosomes includes two technical duplicates. **(B)** Volcano plot depicting the top differentially regulated proteins in AD versus control synaptosomes. **(C)** Correlation analysis of the expression of the top differentially regulated proteins in AD vs. control synaptosome shows a significant R value. **(D)** KEGG pathway analysis of the top upregulated proteins shows significant fold enrichment and protein counts in human diseases and biological pathways. **(E)** KEGG pathway analysis of top downregulated proteins shows significant fold enrichment and protein count in human diseases and biological pathways.

### Status of top synaptosome proteins in Alzheimer’s progression

Based on the biological functions of identified proteins, we validated the levels of the top four deregulated proteins – downregulated (GPI and UQCRC1) and upregulated (TIMM50 and VAT1L) in AD progression.

Immunostaining analyses were performed for the GPI, UQCRC1, TIMM50, and VAT1L proteins on AD postmortem brain samples with different Braak stages and control brains. The results showed that the level of two downregulated proteins, GPI (Glucose Phosphate Isomerase) and UQCRC1, were changed in AD. However, significant downregulation was found in GPI levels with AD severity (Braak Stages V/VI) (**Figure 4**).

**Figure 4.**
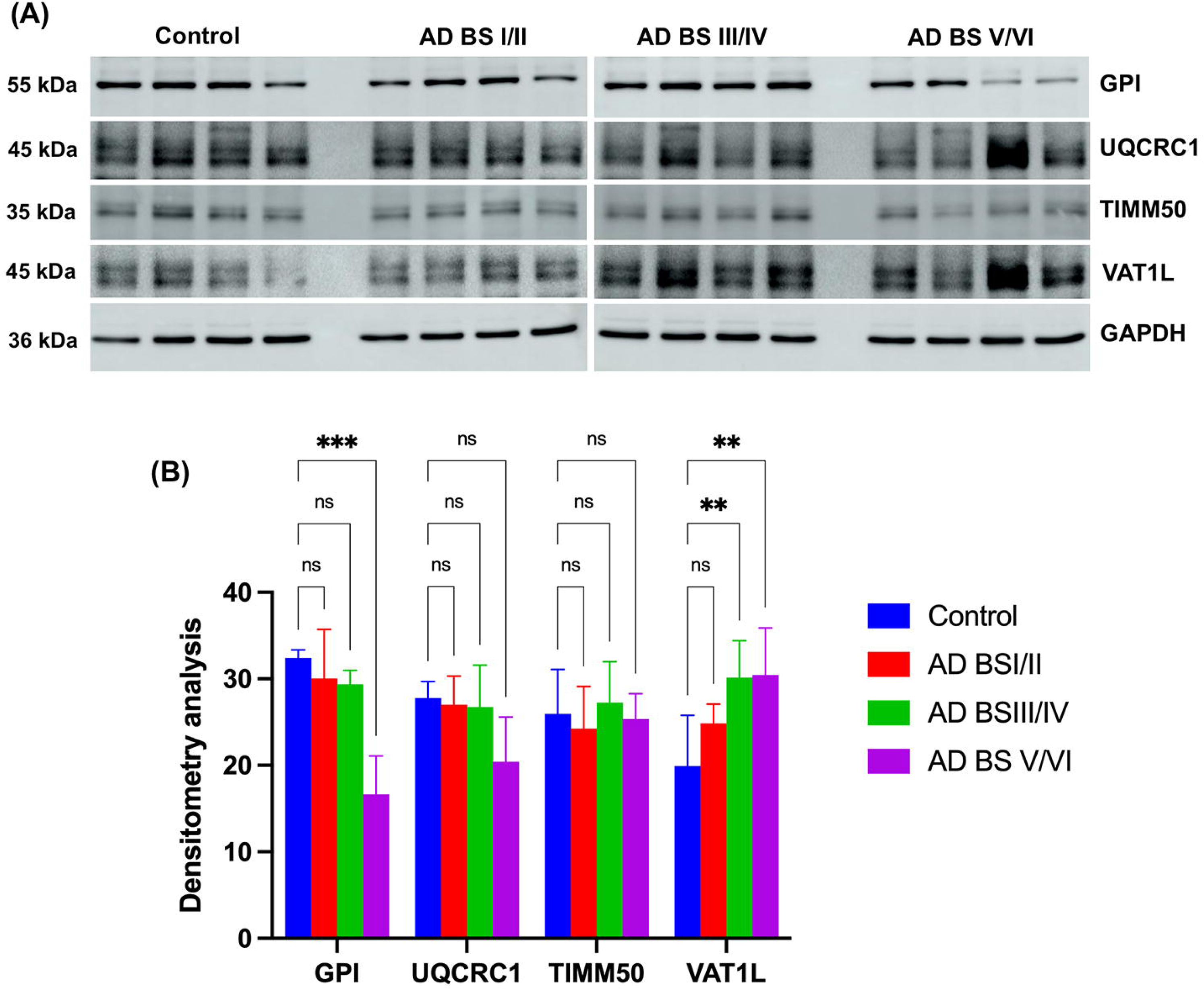
Immunoblotting analysis of top deregulated proteins in AD postmortem brain samples. **(A)** Western blots for GPI, UQCRC1, TIMM50, and VAT1L proteins in postmortem brains from control and AD brains at Braak stages I/II, III/IV, and V/VI. **(B)** Densitometry analysis of GPI, UQCRC1, TIMM50, and VAT1L protein blots in control and AD postmortem brains with different Braak stages. (**P<0.01, ***P<0.001).

We did not observe any significant change in the level of TIMM50 (Translocase of Inner Mitochondrial Membrane 50). However, the level of another top-upregulated protein, VAT1L (vesicle amine transport 1 like), significantly increased in AD with increased Braak stages (**Figure 4**).

Since immunoblotting was performed using the cortical tissue and not on the isolated synaptosomes, this could explain why all four proteins are not showing changes in AD progression. Nonetheless, our results identified two new synapse-associated proteins (GPI and VAT1L) that significantly changed AD progression.

### Molecular interaction of synapse-specific miRNA-mRNA-Protein in Alzheimer’s disease

After conducting analyses specific to each dataset, we examined the molecular interactions of deregulated miRNA, mRNA, and proteins in individuals with Alzheimer’s disease (AD) compared to controls. We found that the fold expression intensity of miRNAs is inversely related to their target mRNA and proteins (**Figure 5**). Most miRNA expression is low (green) while their target protein expression is higher (red). However, we did not observe a significant change in the mRNA expressions of specific proteins. Nevertheless, the top deregulated miRNAs - miR-122-5p, miR-132-3p, miR-194-5p, miR-212-3p, miR-34c-5p, miR-3200-3p, miR-192-5p, miR-421, and miR-340-5p - could negatively modulate their target mRNA translation and protein expression, as shown in the heatmap of **Figure 5**. Furthermore, the gene network map revealed the interaction of each miRNA with their target proteins that were deregulated in AD synaptosomes. Each miRNA candidate interacted with multiple protein targets (**Figure 6**). This analysis demonstrated that the top deregulated miRNA, mRNA, and protein expression may be interdependent, and changes in miRNA expression may lead to the deregulation of their target proteins.

**Figure 5.**
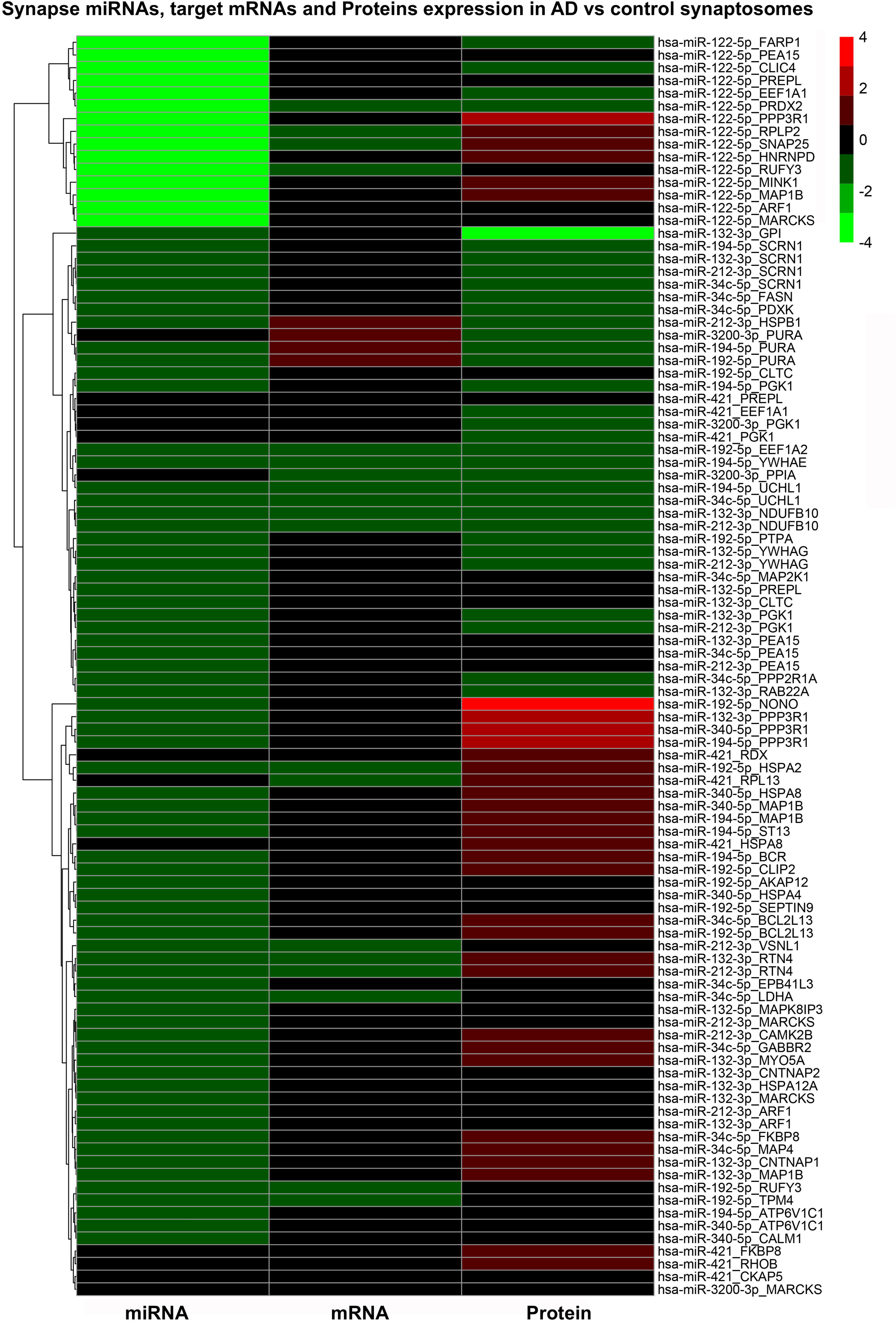
Multi-omics integration analysis of synapse miRNA-mRNA-proteins in AD. Heat map showing the expression levels of miRNAs, their target mRNA, and proteins in AD synaptosomes.

**Figure 6.**
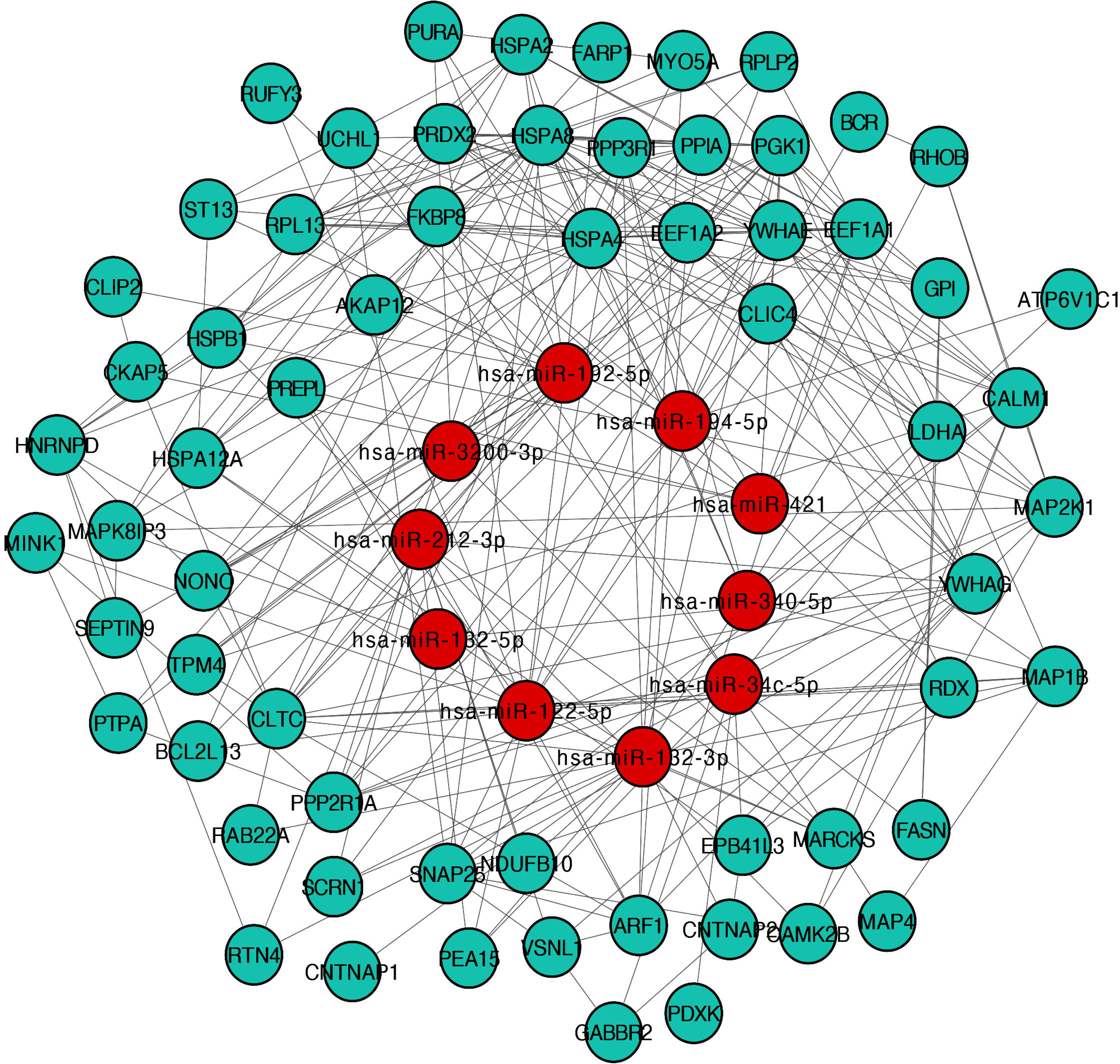
Gene integration analysis of top deregulated miRNAs and their interaction with target proteins in AD synapse.

### Multi-omics integration analysis of synaptosomal miRNA, mRNA, and protein

In addition, to better understand the molecular relationship between synapse miRNA, mRNA, and protein in the same individuals (both with AD and controls), we carried out *D*ata *I*ntegration *A*nalysis for *B*iomarker *D*iscovery using Latent cOmponents (DIABLO). This is a multi-omics integrative analysis aimed at identifying differences between individuals with AD and controls (Singh et al., 2019). For the DIABLO analysis, we selected five control samples and five samples from individuals with severe AD (Braak stage VI).

*A) Circos plot for an integrative framework:* The Circos plot, derived from the DIABLO analysis, reveals intricate relationships between different molecular data types - mRNA, miRNA, and proteins - from AD patients and healthy controls (**Figure 7A**). By integrating these multi-omics data, the plot highlights significant correlations, showcasing how various molecular features interact within the context of AD. Using the first two components (Comp 1-2) allows for a focused representation of the most critical patterns observed in the data, providing insights into potential biomarkers and underlying biological mechanisms of AD. The plot emphasizes features with strong correlations (absolute value of r ≥ 0.7), differentiating positive and negative correlations through distinct connecting lines. For instance, mRNAs such as BAHCC1 and CAMK2N1, miRNAs like hsa-miR-1260a and hsa-miR-124-5p, and proteins including EEF2 and USO1 are shown to have significant interactions. These high-correlation features suggest potential regulatory relationships or co-involvement in disease-related pathways. The differential expression of these features between AD patients and healthy controls further underscores their relevance. For example, miRNAs such as hsa-miR-1260a may regulate mRNA targets implicated in neuronal function or pathology, while proteins like EEF2 and USO1 might play roles in cellular processes disrupted in AD.
*B) Clustered image map representing the multi-omics molecular signature expression:* The heatmap generated from the DIABLO analysis provides a comprehensive visualization of multi-omics data integration for AD (**Figure 7B**). This figure illustrates the expression levels of various molecular features, including mRNA, miRNA, and protein, across samples from AD patients and controls. The color gradient from blue to red represents the range of expression levels, with blue indicating lower expression and red indicating higher expression. The observed patterns highlight significant molecular differences between AD patients and healthy controls.

**Figure 7.**
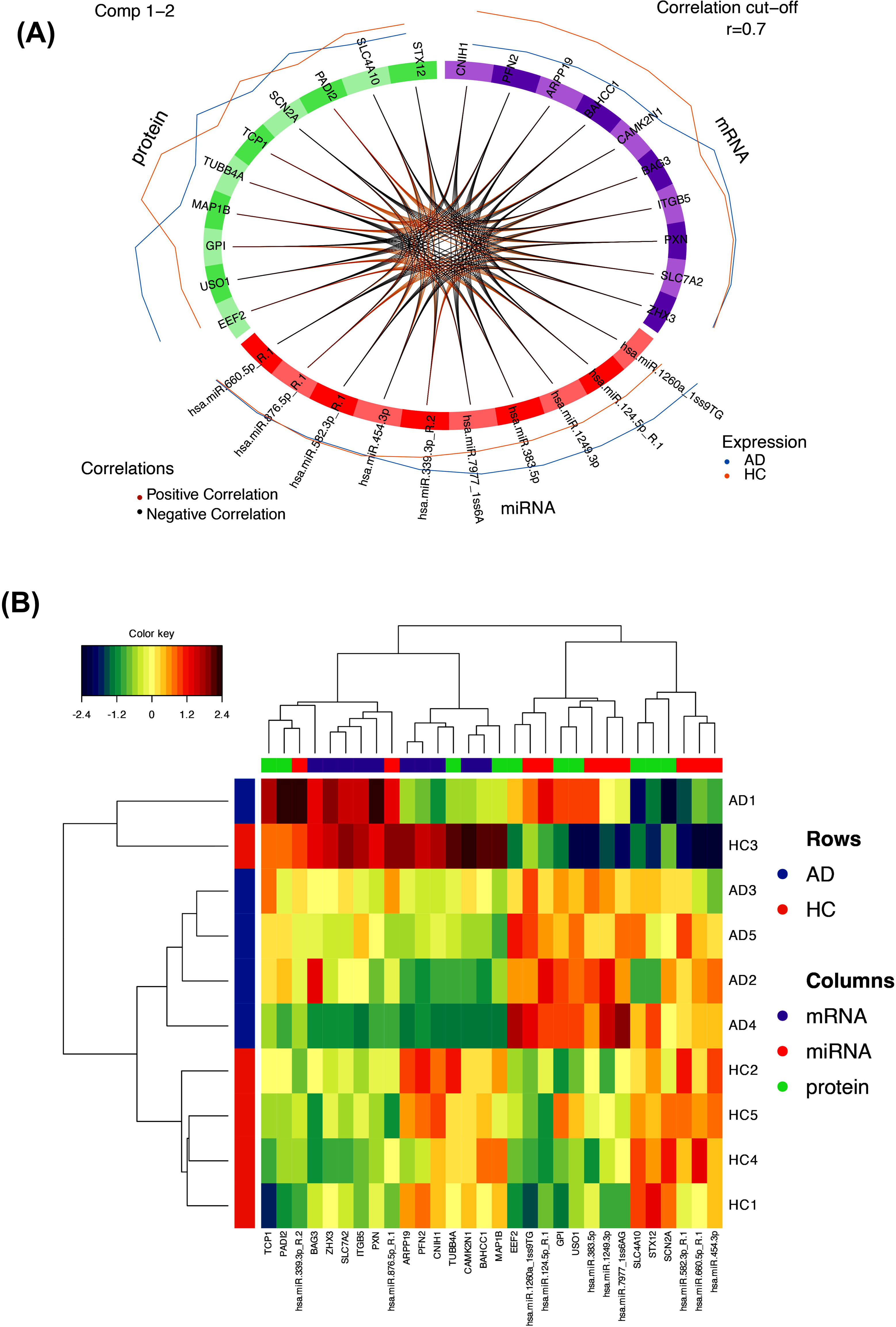
DIABLO analysis of synapse miRNA-mRNA and proteins in AD. **(A)** Circos plot showing the positive and negative correlations between different molecular data types: mRNA, miRNA, and proteins in AD patients and healthy controls. **(B)** The clustered heatmap shows the expression levels of various molecular features, including mRNAs, miRNAs, and proteins, across samples from AD patients and controls.

In the heatmap, mRNAs such as *BAG3*, *ZHX3*, and *CAMK2N1*, miRNAs like hsa-miR-1260a and hsa-miR-124-5p, and proteins including EEF2 and USO1 showed varying expression levels between AD and HC samples. These differential expressions suggest their potential role as biomarkers for AD. For instance, upregulated genes and miRNAs in AD samples (indicated by red shades) could be involved in disrupted pathways. Clustering features with similar expression profiles indicate co-regulation or involvement in related biological processes, further emphasizing their relevance in the disease context.

## Discussion

AD is a synaptic failure disease; therefore, it is of utmost importance to understand the in-depth molecular biology behind the synapse function and dysfunction in AD. Synapse is a small cargo that retains specific RNAs and proteins that work together for normal synaptic plasticity. So far, multiple genes and proteins have been identified at synapses that play very close roles in neurotransmission and synaptic activity (44). Deregulation of these synaptic genes leads to synapse dysfunction in AD. Other than mRNA and synaptic proteins, several small RNAs (miRNAs) have also been identified in AD synapses that play critical roles (5,20,45). A homeostatic balance among miRNA, mRNA, and protein expression is necessary for a healthy synapse function. It is well known that AD synaptic dysfunction in AD is initiated by multiple factors such as amyloid-ß and p-tau proteins, aging, inflammation, environmental factors, and other genetic and epigenetic factors. However, the exact molecular mechanism of synapse dysfunction in AD is largely unknown. Is it not clear whether the deregulation of specific miRNAs, mRNAs, or proteins causes synaptic dysfunction in AD, or if all three molecular subsets work together and lead to synaptic dysfunction? In the current study, we implemented a multi-omics integrative analysis to understand the status of each molecular subset: miRNAs, mRNAs, and proteins in AD. How each molecular subset changes in AD relative to control situations and what the connections are among each subset.

### Changes in synapse miRNAs in AD

We first analyzed changes in the small RNA transcriptome, specifically miRNA levels in the synaptosome fraction isolated from postmortem brains of individuals with AD and the control group. We found both novel and previously reported miRNAs in AD. Among the most significantly downregulated miRNAs in AD are miR-132 (46), miR-34 (47), and miR-212-3p (48), which have been extensively studied in AD. Another significantly downregulated miRNA, miR-122-5p (more than 6-fold down), is less studied in the context of AD. Recent research has identified the role of miR-122-5p in compromised microglial chemotaxis and reduced restrictions of AD pathology (49). Its higher synapse content further emphasizes its relevance in AD research. Additionally, gene set enrichment analysis revealed the significant involvement of downregulated miRNAs in AD and various pathways related to AD and brain function. On the other hand, the top upregulated miRNA, miR-199a, is involved in AD development by regulating Neuritin expression in APP/PS1 mice (50). Furthermore, miR-451a has recently been explored as a serum biomarker in AD (51), and miR-144 is being investigated in AD pathogenesis as a key modulator of ADAM10 protein (52). However, there is limited information available for miR-1247-5p in the context of AD. Additionally, downregulated miRNAs were found to be involved in AD and synapse-related biological pathways.

### Changes in synapse mRNAs in AD

Similar to miRNAs, we also investigated the changes in the expression of protein-coding genes from the same samples of individuals with AD and controls. As expected, we found several genes whose expression was significantly affected in AD compared to the controls. Most of these deregulated genes had been previously associated with AD, but their functions in relation to AD have been minimally studied. The top-upregulated gene, *SHANK1*, has been linked to neuropsychiatric disorders and cognitive dysfunction in various neurodegenerative diseases (53,54). Variants of the *HIVEP* gene have been reported to be dysregulated in AD (55). Similarly, the downregulated gene *TF* (Transferrin) has been well-studied in AD and is associated with AD risk factors (56). Another gene, *TUBA1A* (Tubulin), is also associated with trafficking defects and impaired motor behavior (57). Our gene set enrichment analysis (**SI Figure 5**) revealed the significant involvement of downregulated genes in brain-related biological pathways such as neuronal function, synaptic function, and AD progression. Many of these genes are involved in AD either directly or indirectly; however, further research is required to understand the roles of these genes in relation to AD.

### Changes in synapse proteins in AD

In a manner similar to miRNAs and mRNAs, our proteomic analysis revealed interesting proteins that were deregulated in AD compared to controls. These proteins have barely been investigated for their role in AD. One of the novel proteins downregulated in AD synapses was GPI (Glucose Phosphate Isomerase). Validation analysis also showed the downregulation of GPI with AD Braak stages. This is the first time GPI proteins have been reported in AD. GPI regulates glucose metabolism, and its deficiency causes fetal hemolytic anemia (58). Since synapse functioning requires regulated glucose metabolism and ATP, it is very important to investigate the role of GPI in synaptic function and AD.

Another downregulated protein was UQCRC1 (Ubiquinol-Cytochrome C Reductase Core Protein 1); however, our validation analysis did not show any significant changes in UQCRC1 levels in AD postmortem brains. Similarly, the top-upregulated novel proteins were TIMM50, NONO, and VAT1L. Based on their biological functions, we validated the levels of TIMM50 and VAT1L in AD postmortem brains, showing significant changes in VAT1L levels. VAT1L has been identified in the brain and is mainly associated with cellular functions related to neuronal maintenance, neurotransmission, and Tau pathology (59). Therefore, further research is needed to understand the role of VAT1L in mitochondrial dysfunction at AD synapses.

### Multi-omics integration analysis reveals an intricate relationship between miRNAs, target mRNA, and proteins

The integration of multi-omics data in AD revealed a connection between deregulated miRNAs and their target proteins. One specific downregulated miRNA, miR-122-5p, is associated with certain upregulated target proteins in AD. This analysis provides an overview of disrupted miRNA-mRNA-protein interactions in AD and suggests strategies to restore their normal function. Integrating multi-omics data using DIABLO provides a powerful approach to understanding the molecular complexity of AD. By identifying highly correlated and differentially expressed mRNAs, miRNAs, and proteins, researchers can identify potential biomarkers for early diagnosis or therapeutic targets. These molecular features may contribute to the development of AD through various mechanisms, including altered gene expression, disrupted signaling pathways, or impaired protein functions. For example, BAHCC1 and CAMK2N1 may be involved in neuroinflammatory responses or synaptic signaling alterations in AD, while specific miRNAs may modulate these processes by targeting relevant mRNAs.

Furthermore, the DIABLO analysis helps in gaining a comprehensive understanding of the disease by linking different types of molecular data. This broader view can lead to discovering new interactions and pathways that may be missed in studies focusing on a single type of molecular data. As a result, the insights obtained from this integrated approach not only improve our understanding of the molecular basis of AD but also open up possibilities for developing therapeutic strategies targeting multiple factors.

The grouping of samples into distinct categories (AD and HC) based on their molecular profiles emphasizes the power of integrating multiple types of molecular data in distinguishing between healthy and diseased states. The ability to differentiate AD patients from healthy individuals based on their molecular signatures highlights the diagnostic potential of these biomarkers. Additionally, the identified mRNAs, miRNAs, and proteins offer insights into the underlying biological mechanisms of AD. For example, the increased expression of specific miRNAs in AD samples may indicate their role in regulating genes involved in neurodegeneration, while differentially expressed proteins could point to disrupted cellular functions.

Overall, the multi-omics data aided in identifying key molecular signatures associated with AD. The insights gained from this analysis enhance our understanding of the molecular basis of the disease and pave the way for developing targeted diagnostic and therapeutic strategies. It is essential to validate these findings in larger and more diverse groups to translate them into clinical applications, ultimately aiming to improve the diagnosis, prognosis, and treatment of AD.

In conclusion, our study identified novel miRNA, mRNA, and protein targets centered on synapses in AD within the same individuals under diseased and normal conditions.

## Acknowledgments

The authors are exceedingly grateful to Prof. Rajkumar Lakshmanaswamy, Chair of the Department of Molecular and Translational Medicine, TTUHSC El Paso, for the immense research support. We thank our lab members, Mrs. Shery Rodriguez and Ms. Gunjan Goyal.

## Author Contributions

Conceptualization and supervision: SK; experimental performance: SK, ER, AH, BS, MMT, and DR; analysis, interpretation, and validation of data: SK, SG, and ER; writing and original draft preparation: SK, ER and DD; review, editing, and finalization of manuscript: SG, ER, and SK. All authors have read and agreed to the published version of the manuscript.

## Data Availability

Data for RNA sequencing is available on NCBI Gene Expression Omnibus (GEO) for the following datasets: mRNA (GSE276756) and miRNA (GSE276898). Datasets for proteomics are available on ProteomeXchange via the PRIDE (EMBL-EBI) database (PXD055784).

