## Supplementary figures and images for "Integrated Multi-Omics Analyses of Synaptosomes Revealed Synapse-Centered Novel Targets in Alzheimer’s Disease"

### SI Fig. 2 Complete heatmap of differentially expressed miRNAs in AD vs control synaptosomes.pdf

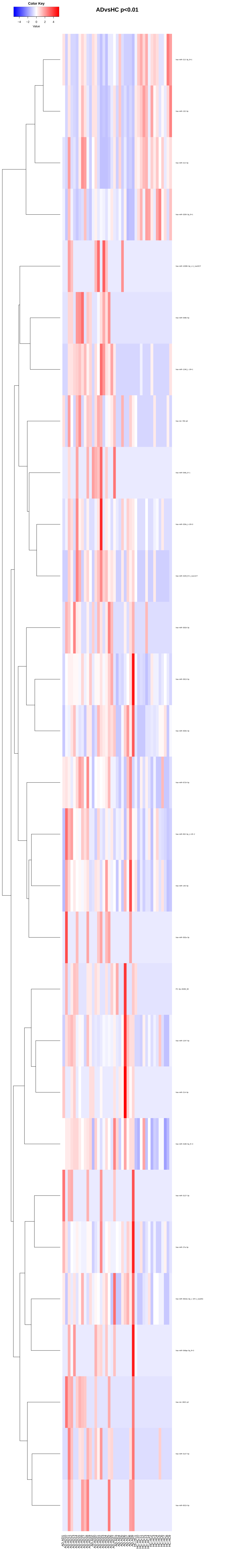

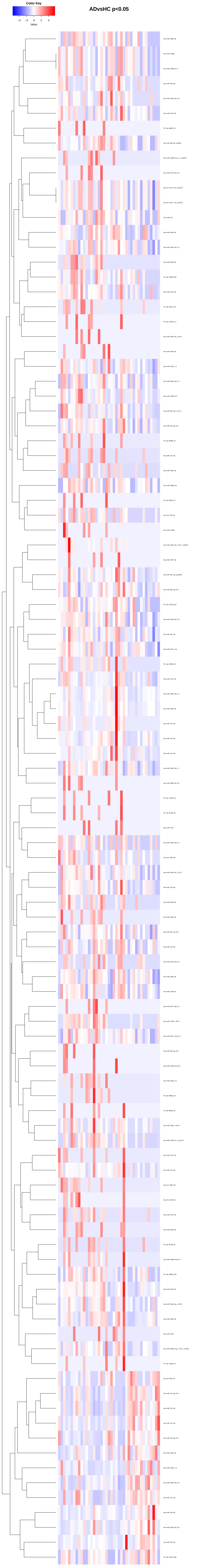

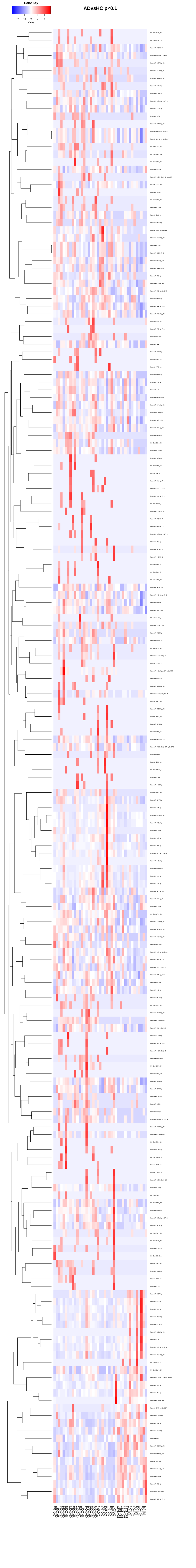

### SI Figure 3

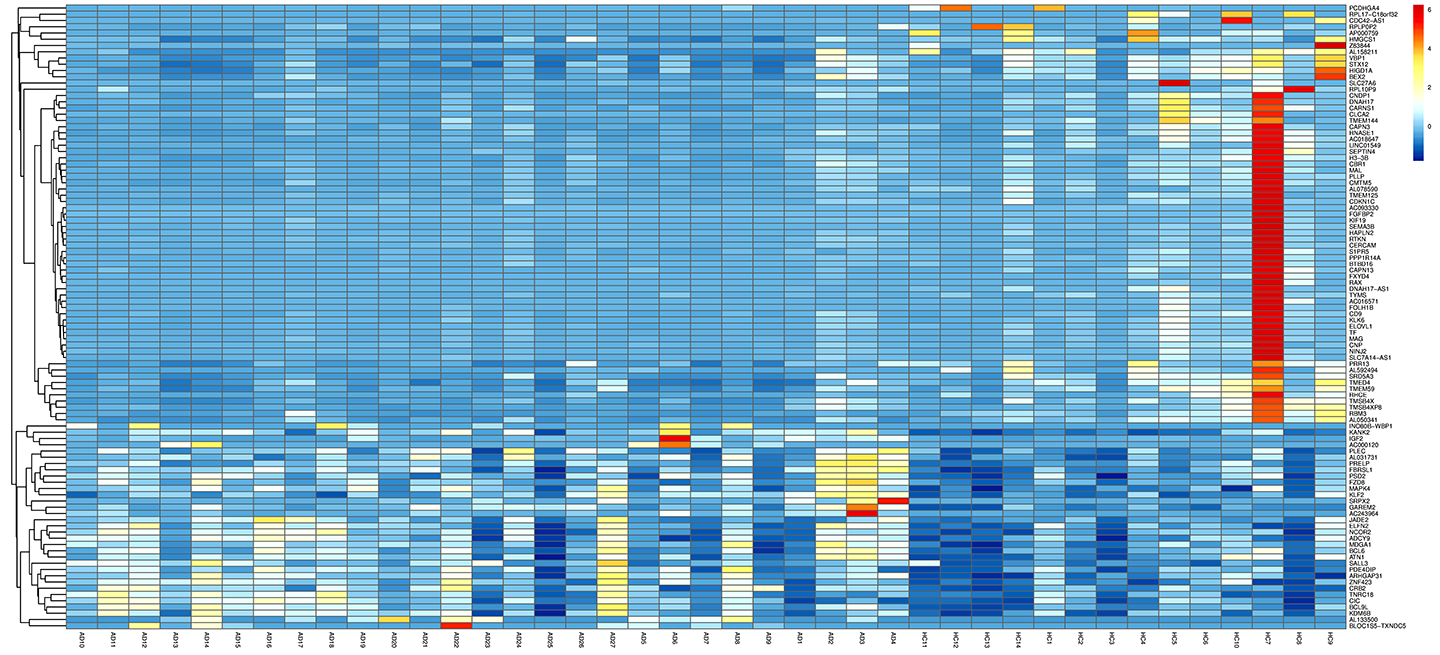

### SI Figure 4

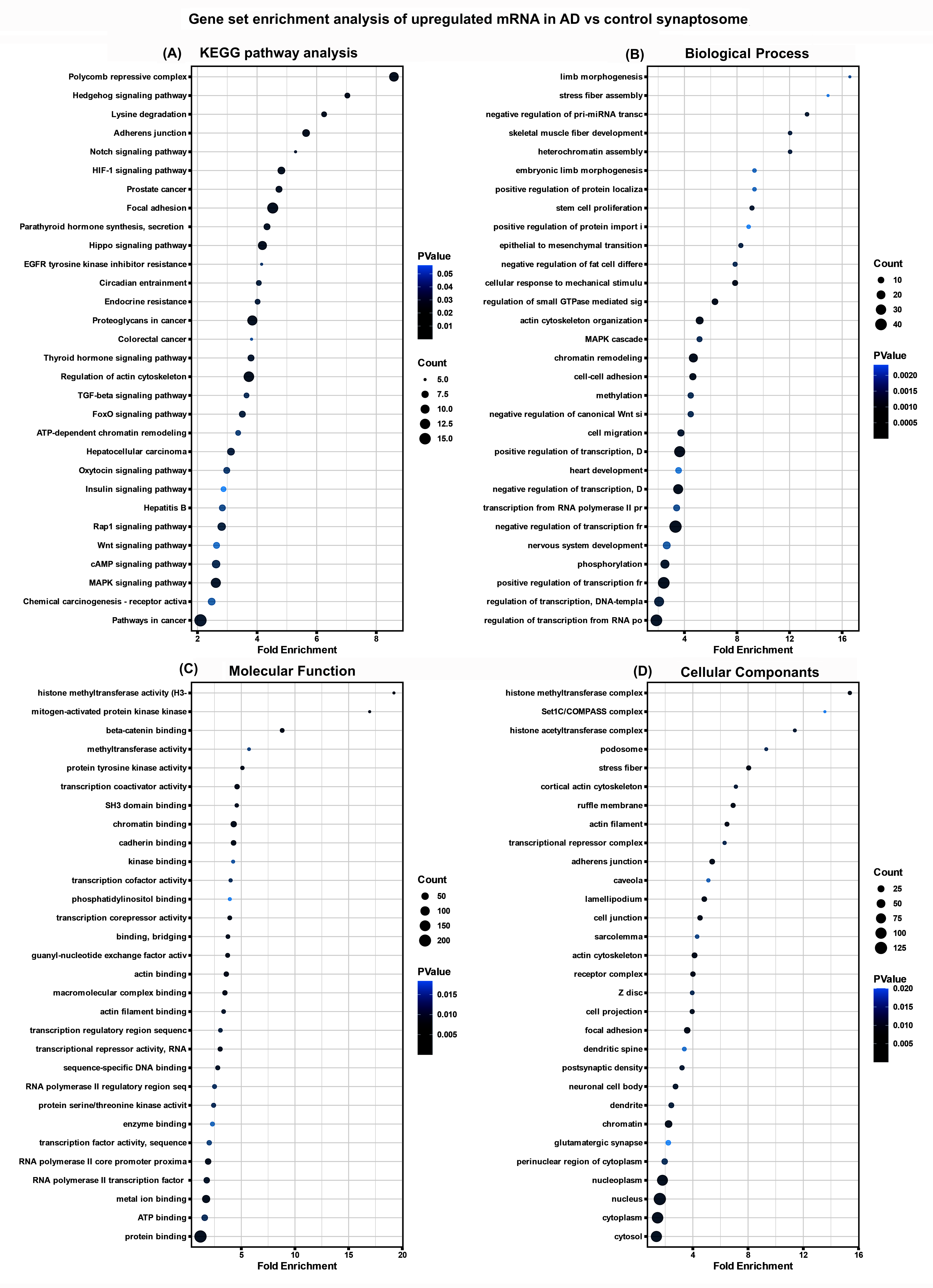

### SI Figure 5

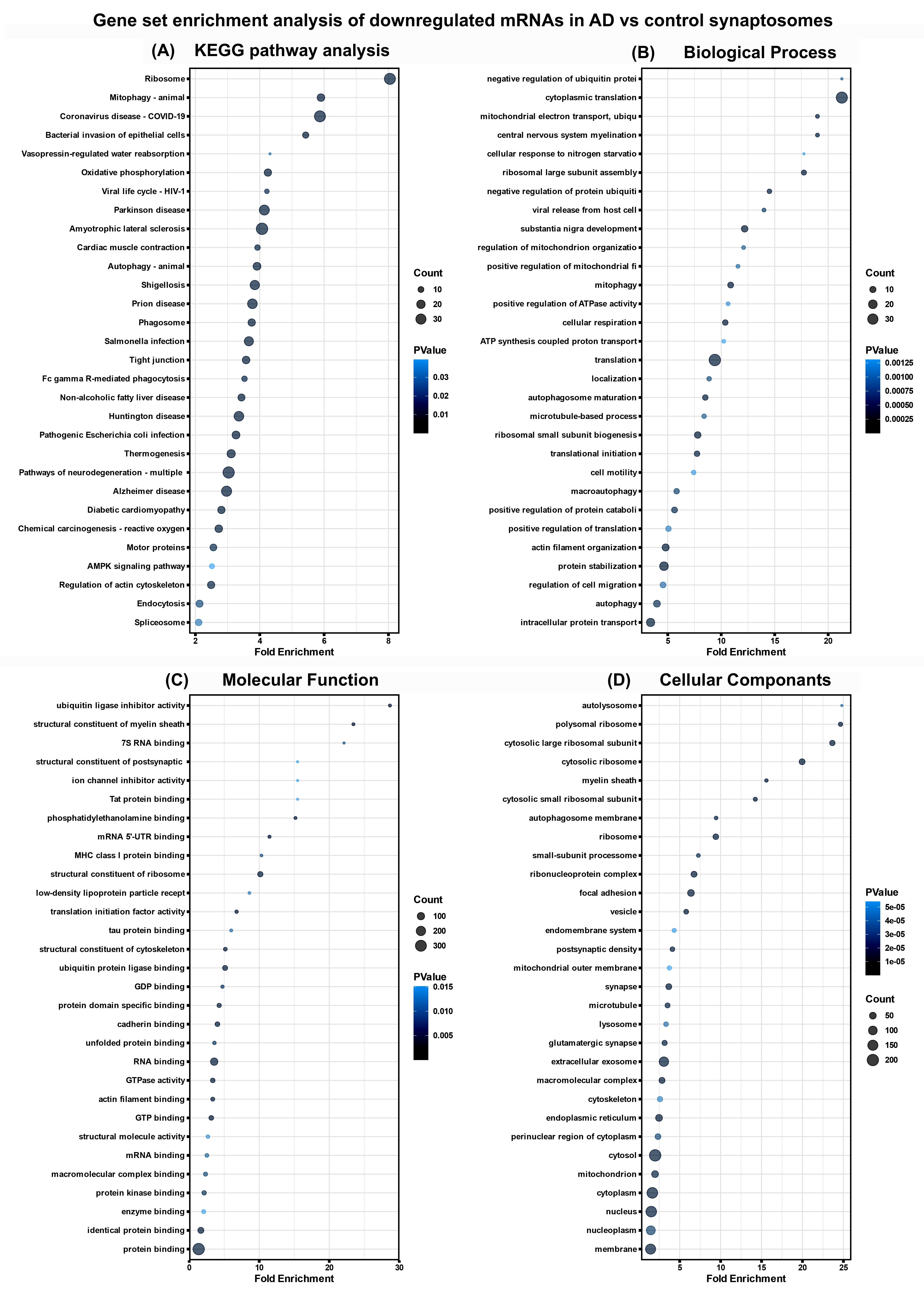

### SI Figure 6

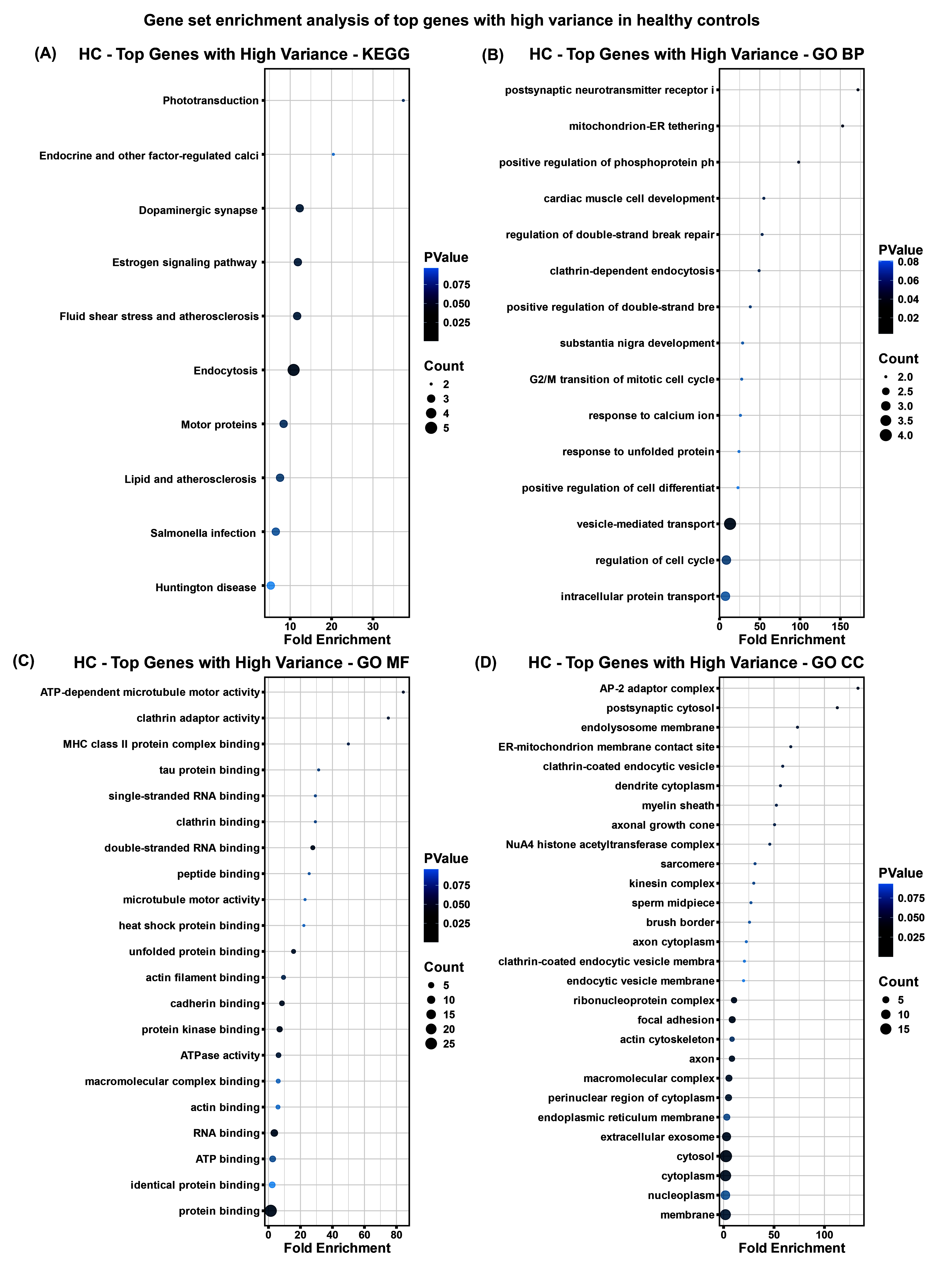

### SI Figure 7

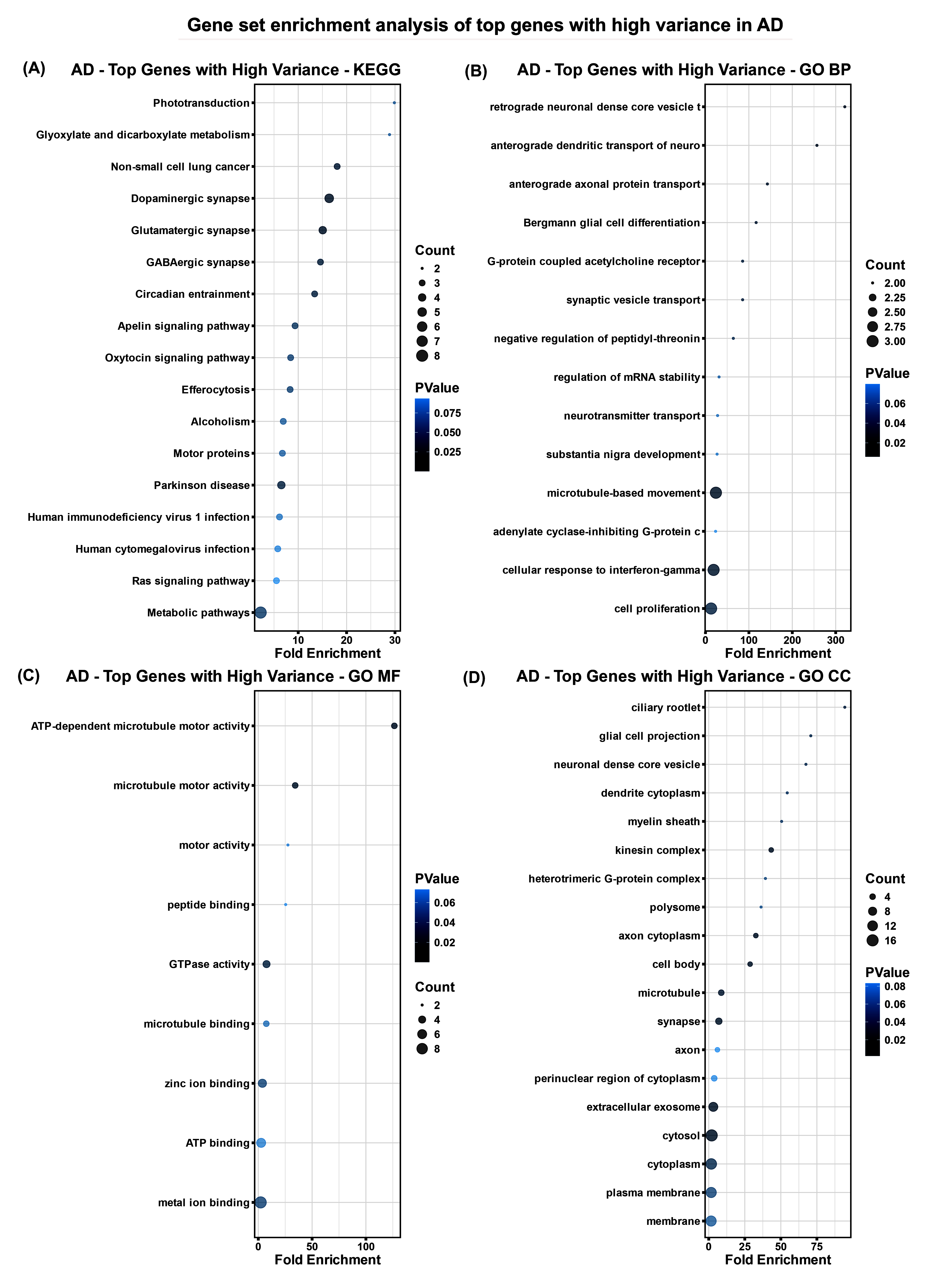
