## Supplementary material for "Integrated Multi-Omics Analyses of Synaptosomes Revealed Synapse-Centered Novel Targets in Alzheimer’s Disease": SI Table 1

| **Table 1- Details of human postmortem brains** | | | | | | | |
| --- | --- | --- | --- | --- | --- | --- | --- |
| **1. Human Brains and Spinal Fluid Resource center, Los Angeles** | | | | | | | |
| **S. No** | **HSB#** | **Age** | **Sex** | **Neuropathology** | **Coronal slab#** | **Structure** | **Autolysis time (hr)** |
| 1 | 4513 | 74 | M | AD | 1 | Broadmann's Area 10 | 15.6 |
| 2 | 4498 | 76 | M | AD | 1 | Broadmann's Area 10 | 12.9 |
| 3 | 4204 | 68 | M | AD | 1 | Broadmann's Area 10 | 11.9 |
| 4 | 4203 | 72 | F | AD | 1 | Broadmann's Area 10 | 20.3 |
| 5 | 4454 | 82 | F | AD | 1 | Broadmann's Area 10 | 9 |
| 6 | 4043 | 80 | F | AD | 1 | Broadmann's Area 10 | 13 |
| 7 | 4382 | 74 | F | AD | 1 | Broadmann's Area 10 | 16.2 |
| 8 | 4617 | 73 | F | AD | 1 | Broadmann's Area 10 | 18.9 |
| 9 | 4718 | 93 | F | AD | 1 | Broadmann's Area 10 | 8.2 |
| 10 | 4608 | 80 | M | AD | 1 | Broadmann's Area 10 | 3.1 |
| 11 | 4752 | 89 | M | AD | 1 | Broadmann's Area 10 | 9 |
| 12 | 4788 | 65 | M | AD | 1 | Broadmann's Area 10 | 7.8 |
| 13 | 4130 | 67 | F | Normal | 1 | Broadmann's Area 10 | 11.8 |
| 14 | 4431 | 68 | F | Normal | 1 | Broadmann's Area 10 | 23.7 |
| 15 | 4660 | 73 | F | Normal | 1 | Broadmann's Area 10 | 18.5 |
| 16 | 5072 | 83 | M | Normal | 1 | Broadmann's Area 10 | 19.5 |
| **2. Brain Endowment Bank University of Miami** | | | | | | | |
| **S.No.** | **Tissue code** | **Age** | **Sex** | **Brain type** | **Race** | **Structure** | **Autolysis time** |
| 1 | HBFR1703 | 69 | F | AD | C | Broadmann's Area 10 | 22 |
| 2 | HBFQ1711 | 77 | M | AD | C | Broadmann's Area 10 | 18 |
| 3 | HBJG1710 | 79 | M | AD | C | Broadmann's Area 10 | 23.8 |
| 4 | HBDA1704 | 80 | M | AD | C | Broadmann's Area 10 | 22.1 |
| 5 | HCTYN1713 | 80 | F | AD | C | Broadmann's Area 10 | 6.5 |
| 6 | HBDI1710 | 85 | F | AD | C | Broadmann's Area 10 | 8 |
| 7 | HBEM1701 | 86 | M | AD | C | Broadmann's Area 10 | 15.5 |
| 8 | HBIP1701 | 90 | F | AD | C | Broadmann's Area 10 | 22.1 |
| 9 | HBCG1703 | 90 | F | AD | C | Broadmann's Area 10 | 8.5 |
| 10 | HCTZX1702 | 95 | M | AD | C | Broadmann's Area 10 | 19.8 |
| 11 | HCT15HAO1713 | 70 | M | Control | C | Broadmann's Area 10 | 12.7 |
| 12 | HCTZZC1711 | 82 | F | Control | C | Broadmann's Area 10 | 14.2 |
| 13 | HCT15HBC1709 | 83 | M | Control | C | Broadmann's Area 10 | 25 |
| 14 | HCTZZT1702 | 84 | M | Control | C | Broadmann's Area 10 | 15.5 |
| 15 | HCT15HBU1704 | 91 | F | Control | C | Broadmann's Area 10 | 18.7 |
| **3. Mount Sinai NIH Brain and Tissue Repository** | | | | | | | |
| **S. No.** | **Barcode** | **Age** | **Sex** | **DX** | **Race** | **Brain region** | **PMI** |
| 1 | 77423 | 79 | F | AD | W | Broadmann's Area 10 | 6.50 |
| 2 | 77424 | 69 | M | AD | W | Broadmann's Area 10 | 5.42 |
| 3 | 77425 | 75 | M | AD | W | Broadmann's Area 10 | 8.00 |
| 4 | 77426 | 94 | F | AD | W | Broadmann's Area 10 | 4.33 |
| 5 | 77427 | 82 | M | AD | W | Broadmann's Area 10 | 20.67 |
| 6 | 77428 | 65 | M | NL | H | Broadmann's Area 10 | 3.83 |
| 7 | 77431 | 103 | F | NL | W | Broadmann's Area 10 | 3.83 |
| 8 | 77433 | 75 | M | NL | B | Broadmann's Area 10 | 5.00 |
| 9 | 77436 | 93 | M | NL | W | Broadmann's Area 10 | 4.17 |
| 10 | 77437 | 84 | F | NL | W | Broadmann's Area 10 | 5.48 |
