## Supplementary material for "Integrated Multi-Omics Analyses of Synaptosomes Revealed Synapse-Centered Novel Targets in Alzheimer’s Disease": SI Table 2

**Supplementary Information**

**SI Table 2. Summary of antibody dilutions and conditions used in the immunoblotting analysis**

| Marker(s) | Primary Antibody and Dilution(s)  (4^°^C, overnight) | Purchased from Company, City & State | Secondary Antibody, Dilution(s)  (Room temperature, 2 h) | Purchased from Company, City & State |
| --- | --- | --- | --- | --- |
| GPI  (15171-1-AP) | Rabbit polyclonal  1:1000 | Proteintech  Rosemont, IL | Goat anti-rabbit IgG HRP 1:10,000  (A9169-2 mL) | Millipore Sigma  Burlington, MA |
| UQCRC1  (21705-1-AP) | Rabbit polyclonal 1:1000 | Proteintech  Rosemont, IL | Goat anti-rabbit IgG HRP 1:10,000  (A9169-2 mL) | Millipore Sigma  Burlington, MA |
| TIMM50  (22229-1-AP) | Rabbit polyclonal 1:1000 | Proteintech  Rosemont, IL | Goat anti-rabbit IgG HRP 1:10,000  (A9169-2 mL) | Millipore Sigma  Burlington, MA |
| VAT1L  (**PA5-98934**) | Rabbit polyclonal  1:1000 | Thermo Fisher Scientific  Waltham, MA | Goat anti-rabbit IgG HRP 1:10,000  (A9169-2 mL) | Millipore Sigma  Burlington, MA |
| GAPDH  (14C10  ) | Rabbit monoclonal  1:800 | Cell Signaling  Danvers, MA | Goat anti-rabbit IgG HRP 1:10,000  (A9169-2 mL) | Millipore Sigma  Burlington, MA |
